# Nitrogen uptake and macronutrients distribution in mango trees (cv. Keitt) under three N fertigation treatments

**DOI:** 10.1101/2021.07.13.452282

**Authors:** A. Silber, T. Goldberg, O. Shapira, U. Hochberg

**Affiliations:** Institute of Soil, Water and Environmental Sciences, Agricultural Research Organization, the Volcani Center, P.O. Box 15159, Rishon LeZion 7505101, Israel; Department of Food Sciences, Faculty of Sciences and Technology, Tel-Hai College, M.P. Upper Galilee 1221000, Israel; Fruit Storage Research Laboratory, MIGAL – Galilee Technology Center, P.O. Box 831, Kiryat-Shmona 1101600 Israel; Newe Ya’ar Research Center, Agricultural Research Organization, the Volcani Center, P.O. Box 1021, Ramat Yishay 30095, Israel

**Keywords:** Crop load, *Mangifera indica*, fruit:leaf ratio, nitrogen uptake, nitrogen balance, titrable acidity, TSS

## Abstract

We assessed the effects of N concentration in the irrigation water on nutrient uptake and distribution in leaves and fruit of mango cv. Keitt grown in a lysimeter for four years. We applied three treatments: N1 – no N fertilization (less than 2 mg/L in the tap water); N2 – 10 mg/L N; N3 – 20 mg/L N.

Deficient N conditions (N1) generated low vegetative yield, high fruit:leaf ratio, high photosynthetic activity, high leaf P and K concentrations, and high sugar content along with low acidity in the fruit. Excess N concentration (N3) induced vegetative growth, and reduced fruit yield and gas-exchange characteristics. The calculated annual nitrogen uptake heavily depended on the nitrogen supply (N1-26 g/tree; N2-196 g/tree; N3- 185 g/tree).

Fruits were the major N sink being 0.82, 0.26 and 0.05 from the total annual N supplied. The N quantities accumulated in N1 fruits during the reproductive season (May-August), were above the N quantities supplied via fertigation, suggesting that N reserve in the vegetative tissues supplied the fruit’s high N demand.

The finding shows the importance of adequate nitrogen supply to mango trees and the dangers of excessive fertilization.

## 1. Introduction

Mango (*Mangifera indica* L.), called the ‘king of fruit’, is an evergreen tree native to southern Asia. Today, it is cultivated in frost-free tropical and warmer subtropical areas worldwide. Most mango orchards are located in non-irrigated rainy tropical areas, whereas lack of knowledge regarding water uptake and nutrient demands restricts mango productivity under Mediterranean or subtropical climates (Durán Zuazo et al., 2011, 2019).

In principle, fertilization management aims to replace or renew the nutritional elements removed by the crop. Intensive fertilization is required to minimize nutrient deficiencies that might induce alternate bearing or stoichiometric stress, which restrict mango productivity. N is an essential nutrient that may affect many parameters of mango tree productivity, such as: vegetative growth (Raj and Rao, 2006; Stassen and van Vuuren, 1997), alternate bearing (El-Motaium et al., 2019), photosynthesis (Urban et al., 2008), quality of shoot bearing and panicles (Singh et al., 1991), embryo abortion (Singh, 2005), fruit color (Bally, 2007; McKenzie, 1994; Nguyen et al., 2004), and anthracnose disease (Nguyen et al., 2004). Deficient N conditions may depress the vegetative and reproductive development of mango trees, whereas excess N may be harmful because excess vegetative growth impairs floral differentiation (Davenport, 2003, 2009; Pinto et al., 2007; Ramírez and Davenport, 2010). The delicate threshold of Mango’s nitrogen requirements for optimal production is yet to be elucidated.

Leaf-N status is frequently used for fertilization-management decisions in the mango industry (Hundal et al., 2005; Raghupathi et al., 2005; Raj and Rao, 2006; Pinto et al., 2007; Kumar et al., 2015; Salazar-Garcia et al., 2018). However, leaves may remain attached for several years on mango trees (Pinto et al., 2007), and the N-fertilization thresholds may vary with factors such as seasonal aspects (Urban et al., 2006), leaf age, rootstock, physiological and phenological status, distance from the inflorescences or fruit (Urban et al., 2004b; Pinto et al., 2007) and fruit:leaf ratio (Urban and Léchaudel, 2005). Although leaf analysis might be useful (Ramirez and Davenport, 2010), it is not sufficient for optimal management of mango orchards, as it provides an empirical value, and the transformation from concentration (gram N per gram dry weight [DW]) to quantity (gram N per tree or hectare) is not straightforward. To date, flushing lysimeter is the only method to understand trees’ nitrogen balance and its response to various nitrogen availabilities.

In the current study, we investigated the response of mango trees (cv. Keitt) grown in lysimeters to different N-fertilization regimes. Although the root volume and water-stress sensitivity of a lysimeter-grown tree differ from those of a field-grown tree (Goldhamer et al., 1999), this setup provides a major advantage, as it enables direct measurement of plant water and nutrient uptake at high temporal resolution during successive growth stages. The general objective of this research was to assess the effects of fertigation N concentrations on nutrient uptake and distribution in leaves and fruits of ‘Keitt’ mango.

## 2. Materials and methods

### 2.1. General information and site characteristics

The experiments were carried out at the Zemach Experimental Station, located in the northern part of the Jordan Valley, Israel (32° 42′ 09″ N, 35° 35′ 07″ E), near the Jordan River outlet from the Sea of Galilee. The climate is Mediterranean, with mild, wet winters followed by dry, hot summers. Values of global solar radiation, air temperature, vapor pressure deficit (VPD), and evapotranspiration are available in: (https://ims.data.gov.il/ims/1). The rainfall season in the region is between November to April, and annual precipitation at Zemach totaled 245, 380, 542, and 487 mm in 2016-17, 2017-18, 2018-19, and 2019-20 respectively. After 2 years of growth in a nursery, mango trees (*Mangifera indica* L. cv. Keitt) grafted on 13/1 rootstock were planted in May 2014 in 1000-L plastic containers. A plastic sheet covered the containers to prevent evaporation. A volume of 50 L of coarse tuff (volcanic material) was placed at the bottom of the container to ensure proper drainage. The remaining volume was filled with 0–8 mm tuff, a volcanic pyroclastic material that has negligible interaction with the mineral content of the irrigation solution. The drainage from the containers was collected using PVC pipes and conducted to a deep ditch for manual drainage measurements. Tree spacing was similar to that in commercial orchards (3 × 6 m, 555 trees/ha).

### 2.2. Experimental design

The experimental design comprised three treatments, each composed of eight randomized trees. The treatments were:

N1 – no N fertilization (less than 2 mg/L in the tap water)
N2 – 10 mg/L N; equal amounts of NO_3_ and NH_4_
N3 – 20 mg/L N; equal amounts of NO_3_ and NH_4_

All treatments were fertilized with P, K, and micronutrients as described in section 2.3.

The different treatments were begun in November 2015. The irrigation system consisted of 10 pressure-compensated drippers (1.6 L/h, Netafim Inc., Tel Aviv, Israel) per tree, installed around the trunk in each lysimeter. To prevent water shortage, irrigation was started before sunrise and stopped in the late afternoon, applied every 1–1.5 h for 40–50 min using an irrigation controller. All treatments received the same daily amount of water. Irrigation frequency varied from 8 to 12 daily events. The irrigation amount exceeded the tree evapotranspiration to allow for a leaching fraction (ratio of drainage to irrigation amounts) above 0.3, so that the electrical conductivity (EC) of the draining solution could be kept below 1.5 dS/m to prevent salt stress. All of the horticultural practices were performed uniformly according to the recommendations of the Israeli Extension Service. The first yield was harvested at the end of August 2017 and the following yields (2018, 2019, and 2020) were harvested in the same period. Note that each harvest represents the growth season from September (after the preceding harvest) to August.

### 2.3. Fertigation solutions and plant analyses

Samples of the fertigation solutions were periodically collected and analyzed for pH, EC, and nutrient concentrations. The pH and EC of the tap water were 7.7 ± 0.2 and 1.1 ± 0.1 dS/m, respectively. The P and K concentrations in the irrigation solution were 10 and 40 mg/L, respectively. The nutrient solutions were prepared from commercial fertilizers [(NH_4_)_2_SO_4_, NH_4_NO_3_, KNO_3_, KCl and H_3_PO_4_] and tap water containing (mg/L): 0–2 N, 0.1–0.3 P, 9–10 K, 70–80 Ca, 25–30 Mg, 120–140 Na, and 150–190 Cl. Micronutrient concentrations (mg/L) were 0.3 Zn, 0.6 Mn, 1.0 Fe, 0.04 Cu, 0.4 B, and 0.03 Mo, all EDTA-based.

The yields (2017–2020) were harvested at the end of August. Accordingly, we considered the growth season to be from September (after the preceding harvest) to August. All the fruits were collected from all the trees and immediately weighed.

In 2020, the total canopy area was measured after harvest for four trees per treatment by collecting all their leaves and determining their dry weight (DW). DW was converted to the entire canopy area by multiplying it with the leaf mass per area. Leaf mass per area was measured for every tree on 20 leaves. The area of each leaf was measured with a special scanner (leaf area meter, Delta-T Devices, Cambridge, UK) before it was dried for 72 h in a 60 °C oven and weighed. The maximal leaf area index (LAI_max_) was measured near the trunk using a canopy analysis system (SunScan model SS1-159 R3-BF3; Delta-T Devices).

The experimental setup included daily measurements of irrigation and leachate amounts, in addition to analyses of N concentration in the irrigation and leachate solutions sampled every two weeks, which allowed estimating N uptake by the trees. The calculated N quantity taken up from the solution (N-uptake) during 2020 was calculated as follows:

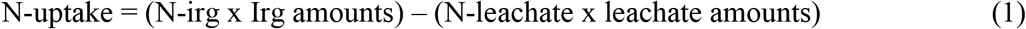

where N-irg = N concentration in the irrigation water; Irg amounts = irrigation quantity (L/tree); N-leachate = N concentration in the leachate; leachate amounts = leachate quantity (L/tree).

Leaves from the concurrent year (formed after the preceding harvest) with the greenest color at a two- to four-node distance from the fruit/flower and reproductive organs (fruitlet and fruit) were sampled from four replicates per treatment every two weeks. The plant material was washed with distilled water, dried in a ventilated oven at 60 °C, and stored pending chemical analysis. The sampled fruit were separated for chemical analyses of peel, seed and mesocarp (pulp). The dry tissue was ground to pass a 20-mesh sieve, and 100-mg samples were wet ashed with H_2_SO_4_^−^–H_2_O_2_ and analyzed for Na, K, organic N, and P. Ashing with HClO_4_–HNO_3_ was used for Ca, Mg, and micronutrient analyses. Element concentrations were determined as already described.

### 2.4. Gas exchange

Gas exchange measurements were taken on 27/05/2020 and 24/08/2020 at midday (12:30) using a commercial gas exchange system (Li-6800; Li-Cor, Inc., Lincoln, NE, USA). Due to the high variability in gas exchange between mango leaves (Urban.Lu *et al*., 2004), we always measured the current year leaves (formed after the last harvest) with the greenest color at a 2-4 nodes distance from the fruit/flower. The conditions were set to PAR of 1600 μmol m^−2^ s^−1^, a flow rate of 1000 μmol s^−1^, boundary layer conductance ~ 3 mol m^−2^ s^−1^, 400 μmol CO_2_ mol^−1^ with ambient humidity and temperature. 3 leaves were measured from each tree and the whole set of 72 leaf measurement was completed within 90 minutes.

### 2.5. Fruit quality, color, sugar, and acidity

The degree of the mango’s skin red color was visually estimated after harvest. Five slices from each replicate were squeezed and centrifuged to obtain clear juice after starch precipitation. Total soluble sugars (TSS) were measured using a digital refractometer (model PR-1, Atago Co., Tokyo, Japan) and were expressed as °Brix (%). Titratable acidity (TA) was assayed by titration of the mango juice with 0.1 N NaOH to pH 8.1 using an automatic titrator (model 719s, Schott Ltd., (St. Gallen) Switzerland). Citric acid is the predominant acid in mango pulp (Léchaudel et al., 2005b) and therefore, the titration data were expressed as percentage of citric acid in the mango juice.

### 2.6. Calculations and statistics

Statistical analyses were carried out with JMP 14 software. All data were analyzed for effects of treatments by means of the general linear model procedure of SAS (SAS Institute, Cary, NC, USA). Differences among means were tested with the standard least-squares model of ANOVA, followed by Tukey HSD pairwise comparison of means. Differences with a probability larger than 95% were considered significant.

## 3. Results

### 3.1. Tree development and yield characteristics

The amount of N-irg significantly affected both vegetative and reproductive yields throughout 2016–2020 (Table 1). Water uptake data for 2019-2020 were used to illustrate the phenological effect on the diurnal water uptake (Table A.1). N1 trees were small, bearing chlorotic leaves that produced low fruit yield (Table 1), however, the minute quantities of N in the tap water (<2 mg/L) still promoted an increase in yield from 17(±2.9) kg/tree in 2017 up to 32(±3.7) kg/tree in 2020. Elevating N-irg to 10 mg/L N (N2) resulted in a significant yield increase (56 kg/tree in 2020) and eliminated the chlorotic leaves. An increase to 20 mg/L N (N3) had a negative effect, reducing mango yield down to 16 kg/tree in 2020 (Table 1).

**Table 1.** Effects of the experimental treatments on selected yield parameters of 2017–2020 seasons.

| Treatment | Fruit<br>(number/tree) | Yield<br>(kg/tree) | Fruit weight<br>(g/fruit) |
| --- | --- | --- | --- |
| 2017 |  |  |  |
| N1 | 35 | 17 | 490 |
| N2 | 89 | 38 | 421 |
| N3 | 71 | 29 | 408 |
| Prob>F<sup>1</sup> | <0.0001 | <0.0001 | <0.0105 |
| LSD<sub>0.05</sub><sup>2</sup> | 22.1 | 10.7 | 82.2 |
| 2018 |  |  |  |
| N1 | 53 | 24 | 459 |
| N2 | 91 | 42 | 458 |
| N3 | 49 | 34 | 711 |
| Prob > F | <0.0001 | 0.0002 | <0.0001 |
| LSD<sub>0.05</sub> | 17.6 | 11.0 | 122.7 |
| 2019 |  |  |  |
| N1 | 63 | 30 | 470 |
| N2 | 146 | 54 | 371 |
| N3 | 98 | 38 | 403 |
| Prob > F | <0.00001 | <0.00001 | 0.0001 |
| LSD<sub>0.05</sub> | 33.9 | 13.0 | 65.3 |
| 2020 |  |  |  |
| N1 | 44 | 32 | 765 |
| N2 | 72 | 56 | 829 |
| N3 | 18 | 16 | 903 |
| Prob > F | <0.0001 | <0.0001 | 0.0495 |
| LSD<sub>0.05</sub> | 29.1 | 15.7 | 136.3 |

<sup>1</sup>: Prob>F for treatments; <sup>2</sup>: LSD<sub>0.05</sub> for treatments

Phenological aspects governed fruit development and accordingly, the rate of DW accumulation of fruit components (peel, pulp, and seed) varied during the reproductive season. The rate of the fruit’s DW accumulation during the early period of fruit development (until 18 June, day of year (DOY) 169) was relatively slow (Fig. 1), but it accelerated later, and the difference between N-fertilization treatments became significant (DOY 204 and 237 represent 23 July and 25 August, respectively). In all treatments and through the four years of the experiment, the average DW proportion of pulp, seed and peel were: 0.67±0.08, 0.21±0.05, and 0.13± x, respectively.

**Fig. 1.**
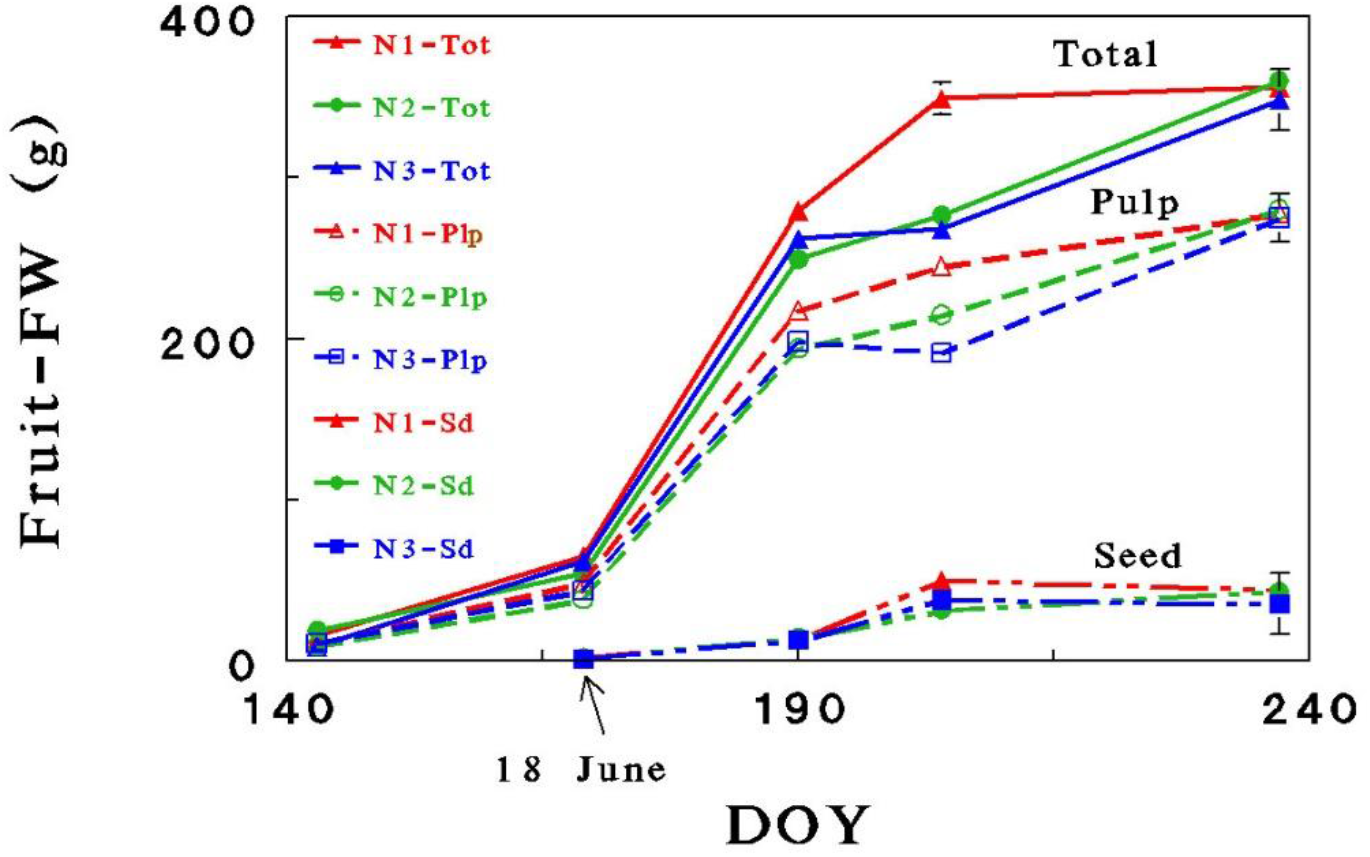
DW accumulation in mango fruit organs: seed (Sd), pulp (Plp) and the total (including seed, pulp and peel) across May-August 2019. DOY is day of the year (20 May is DOY 140). Vertical lines designate standard error values of four trees.

The three N treatments differentially affected the vegetative and the reproductive yields and consequently, induced three different fruit:leaf DW ratio: 3.4, 1.5 and 0.5 for N1, N2 and N3, respectively (Prob > F < 0.0001). Subsequently, three distinct N environments were formed: (i) deficient-N for N1 trees; (ii) adequate-N for N2 trees; and (iii) excessive-N for N3 trees. Single leaf area measured after the 2020 harvest was 64.4(±5.23), 79.2(±1.78) and 87.2(±5.29) cm^2^ for N1, N2 and N3, respectively (Prob > F = 0.0054). Stomatal conductance (*g*_s_) and assimilation rate (An) showed a marked response to the increase in N-irg as well as to the advancing maturation of the fruit (Fig. 2).

**Fig. 2.**
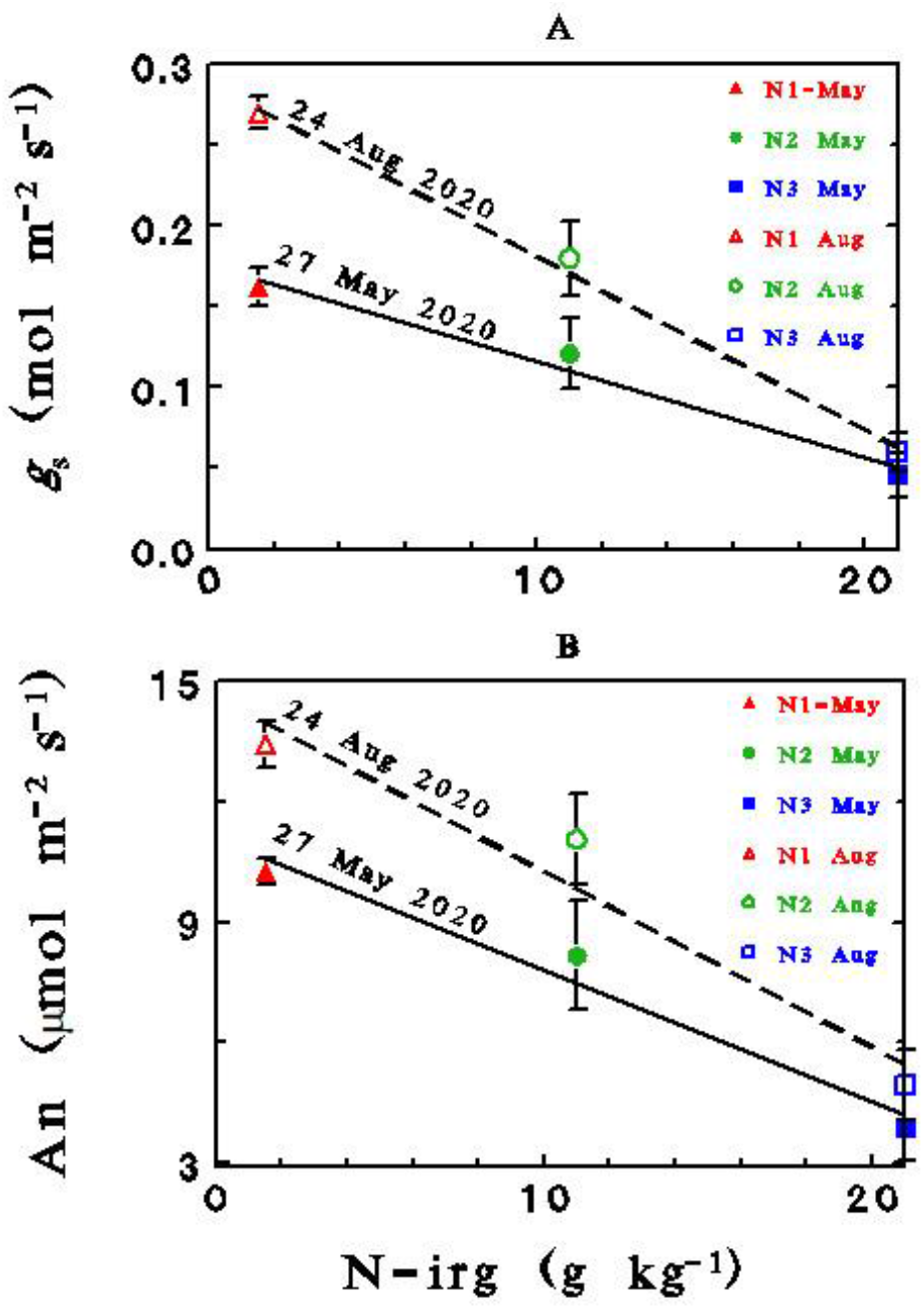
Relationships between N-irg and gas exchange properties on 27 May and 24 August 2020. A: stomatal conductance (*g*_s_), B: photosynthesis (An). Vertical lines designate standard error values of four trees. The fitted equations are: *g*_s_-May – 0.18(0.013)-0.0059(0.0009)*x (r^2^=0.98); *g*_s_-August – 0.29(0.098)-0.011(0.0007)*x (r^2^=0.90); An-May – 11.1(081)-0.327(0.0594)*x (r^2^=0.97); An-August −14.7((1.44)-0.44(0.109*x (r^2^=0.95).

### 3.2. Seasonal fluctuations of N–P–K concentration in leaves and fruit

N-irg, together with mango phenology, determined leaf N-P-K concentrations during the growing season. (Fig. 3). Leaf-N concentrations in N1 trees throughout the growing seasons were above the deficit range of 8 g/kg suggested by Pinto et al. (2007); those of N2 were at the optimum level (12–14 g/kg), and leaf-N concentrations of N3 during the flowering period (March–April) were greatly above the excess threshold. Significant parabolic regression characterized leaf-N concentration of N1 and N2 trees, with a peak in the middle of May (DOY 135), before the stage of rapid fruit development (Fig. 1), whereas the peak for N3 trees was earlier (during the flowering period, March) and higher (Fig. 3). Note that leaf-N concentrations of N3 trees after the flowering period nearly coincided with those of N2 trees. A leaf-N concentration above 14 g/kg may induce vegetative flushes that impair flower induction (Davenport, 2003, 2009) and accordingly, it is probable that the low fruit productivity of N3 trees was directly related to excess N fertilization. Although irrigation P and K concentrations were kept constant for the three treatments, leaf-P and K of N1 trees were significantly higher than in N2 and N3 trees (Fig. 3). Leaf-P concentrations of N2 and N3 trees were at the upper level of 0.8–1.6 g/kg recommended for mango trees by Pinto et al. (2007) and Bally (2009), whereas that of N1 trees markedly surpassed the excess threshold of 2.5 g/kg. Leaf-K of N1 and N2 trees decreased steadily from January to August (harvest), whereas that of N3 trees reached a peak in the middle of May (DOY 135), before the rapid fruit growth (Fig. 1).

**Fig. 3.**
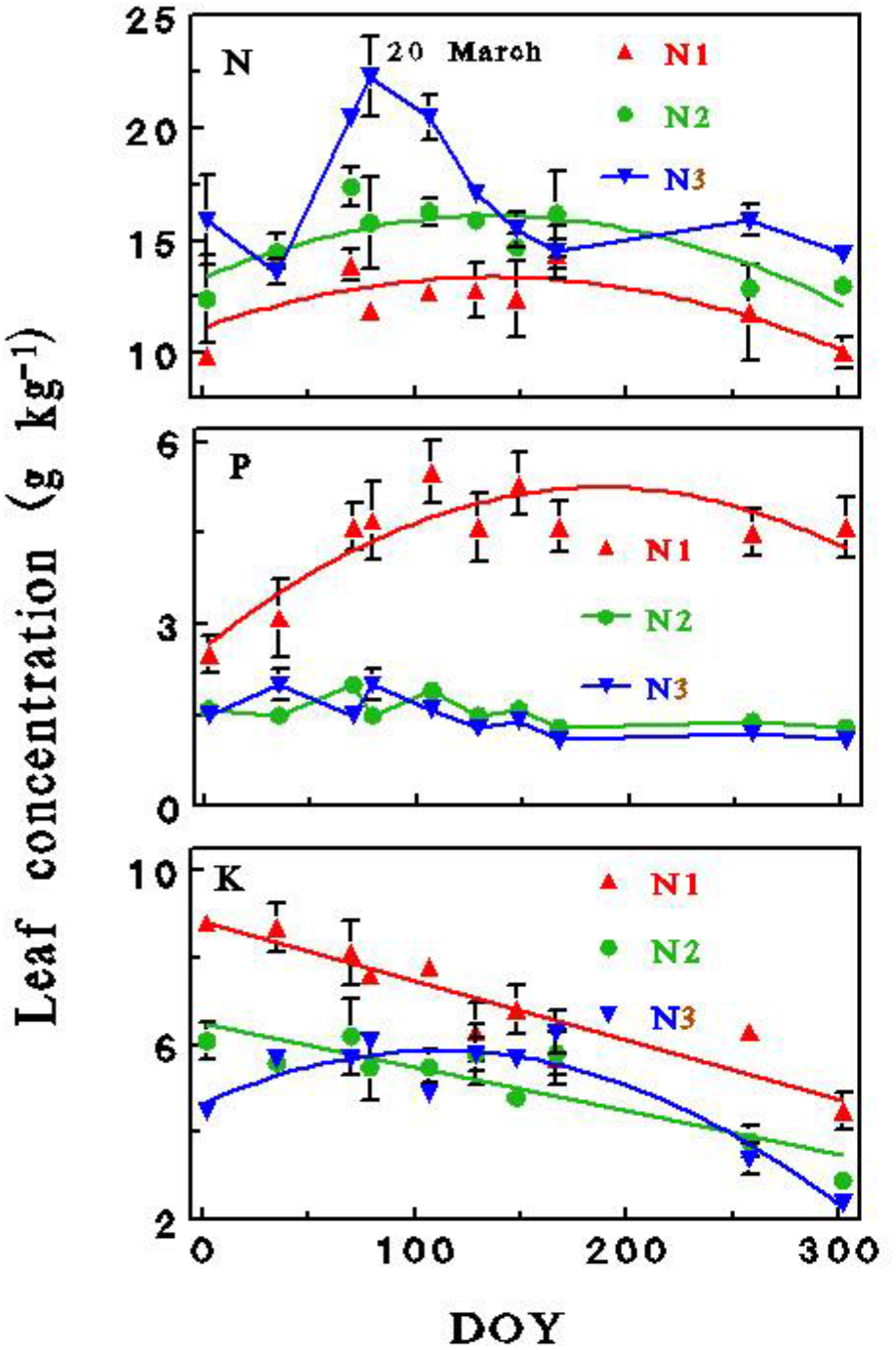
Leaf-N -P and -K concentrations for the different N treatments across 2018-19 years. DOY is day of the year (8 July is DOY 190). The fitted equations are: N – N1: 11.1+0.033*x-0.000012*x^2^ (*r*^2^=0.50); N2: −13.2+0.04*x-0.000043*x^2^ (*r*^2^=0.85); P – N1: 2.6+0.028*x-0.00042*x^2^ (*r*^2^=0.76); K – N1: 8.8-0.013*x (*r*^2^=0.84); N2: 6.5-0.000*x (*r*^2^=0.77); N3: 4.7+0.022*x-0.000099*x^2^ (*r*^2^=0.87). Vertical lines designate standard error values of four trees.

N–P–K concentrations in the pulp and seeds were highest shortly after these organs’ development (DOY 120, representing 30 April) and decreased continuously with increasing organ weight (Fig. 4). Similar to the effect of N-irg on the leaves, N concentrations in the pulp and seeds followed the order: N3 > N2 > N1. Seed-N and P concentrations were significantly higher than in the pulp, and it is logical to assume that immediately after the flowering and fruit-set processes (April), seed-N and P concentrations were even higher, as reported for avocado (Silber et al., 2018). The effect of N-irg treatments on the calculated N–P–K quantities accumulated in a single fruit during 2019 (Fig. 5) was based on the fruit DW and the N–P–K concentrations, presented in Figs. 1 and 4, respectively. Fruit-N, P and K quantities removed by average mango tree (cv. Keitt) bearing, 150 fruit, 400 g each) from the N2 treatment were 70, 10 and 77 g, respectively (Fig. 6). The N-P-K concentrations in fruit organs at 2017-20 harvests are reported in Table 2, A.2 and A.3.

**Fig. 4.**
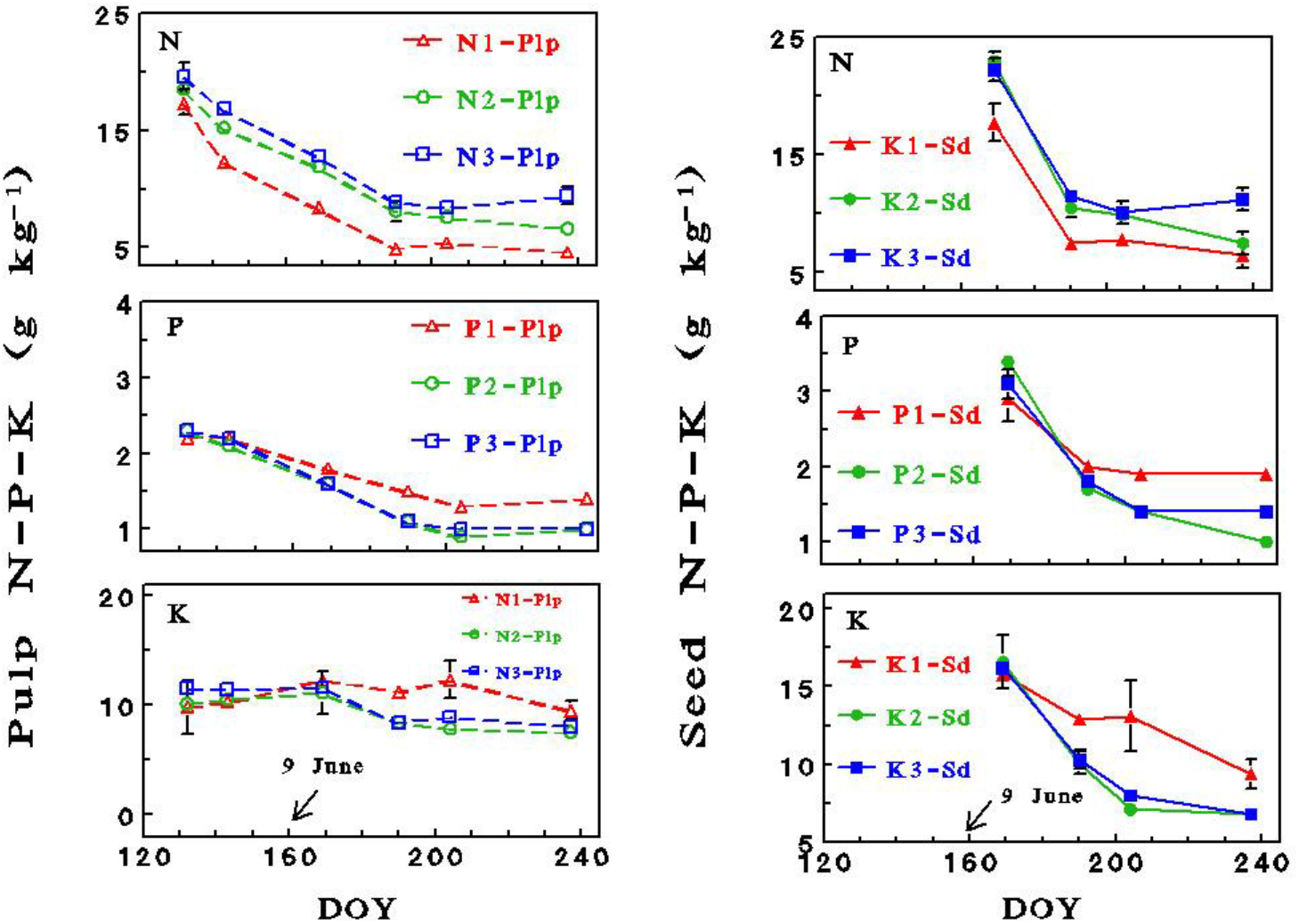
Effect of irrigation-N concentration on N-P-K concentration in seed and pulp of mango fruit across May-August 2018-19 years. DOY is day of the year (9 June is DOY 160). Vertical lines designate standard error values of four trees.

**Fig. 5.**
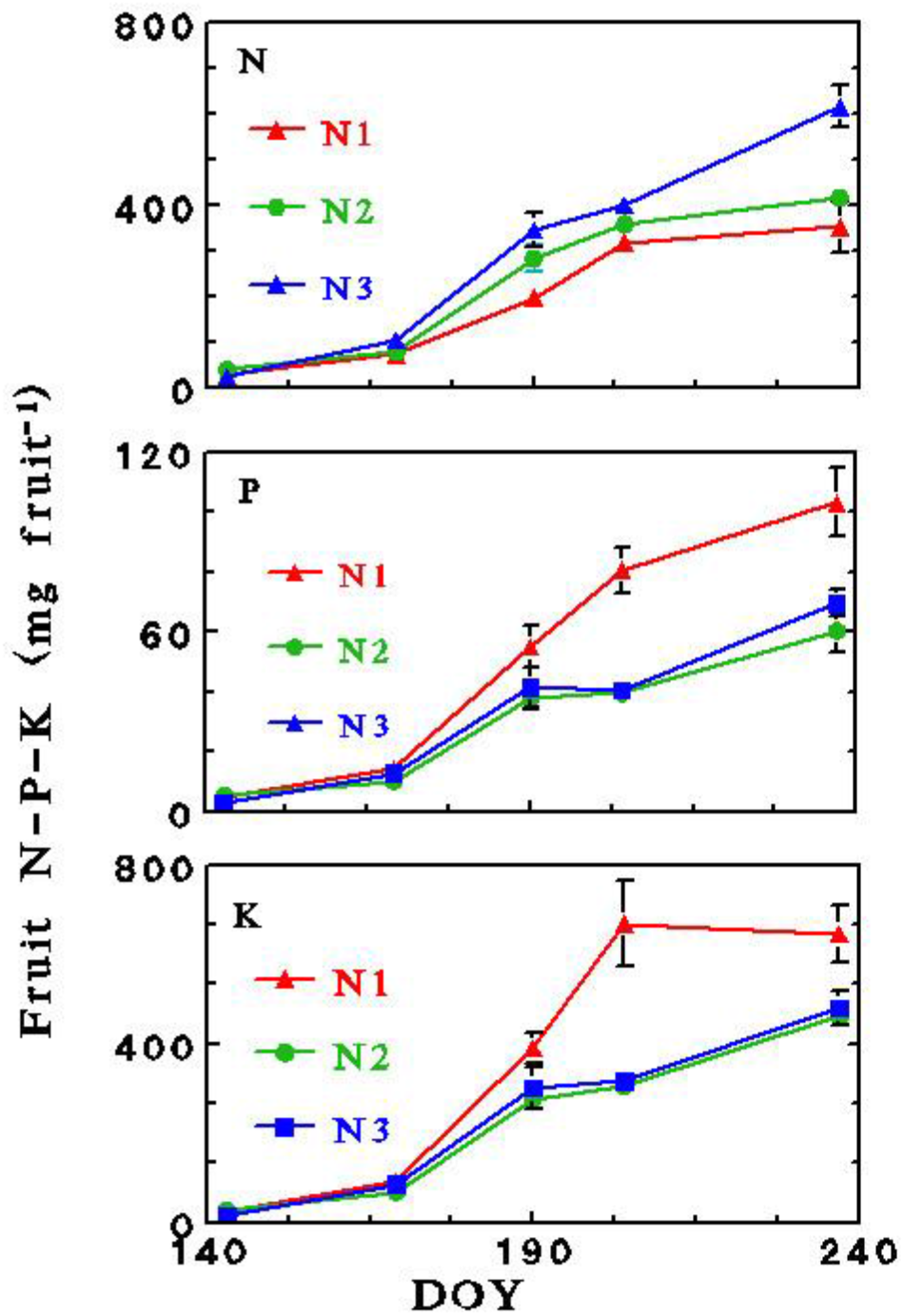
Effect of irrigation-N on N-P-K accumulation in mango fruit (400 g) during May-August 2019. DOY is day of the year (8 July is DOY 190).

**Fig. 6.**
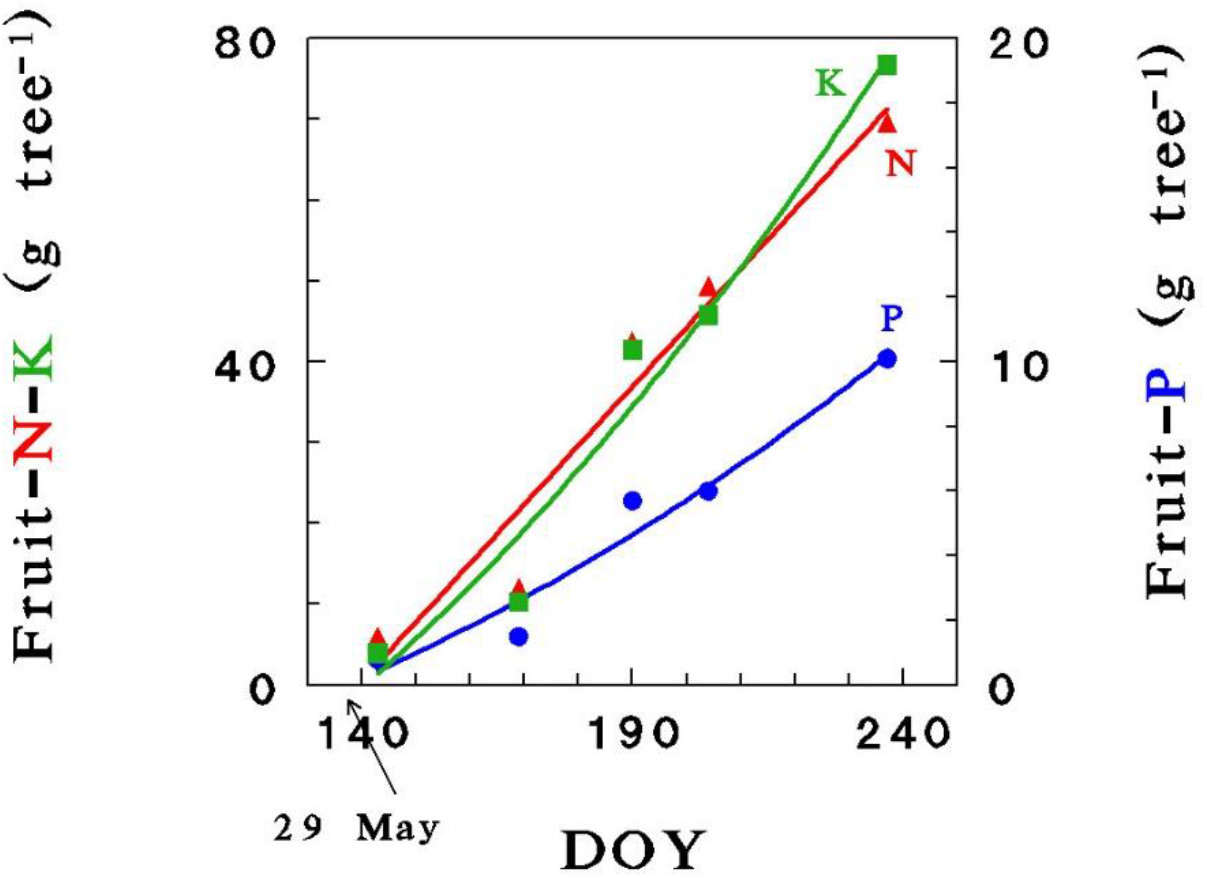
Fruit-N, -P and –K quantities removed by an average high yielding mango (cv. ‘Keitt’) tree (assuming 150 fruit, 400 g each). The figure is based on the 2019 mineral concentration of the fruits, which were similar to the 2017 and 2018, but slightly lower than the 2020 yield (see Table 2, Table S2 and table S3). DOY is day of the year.

**Table 2.**
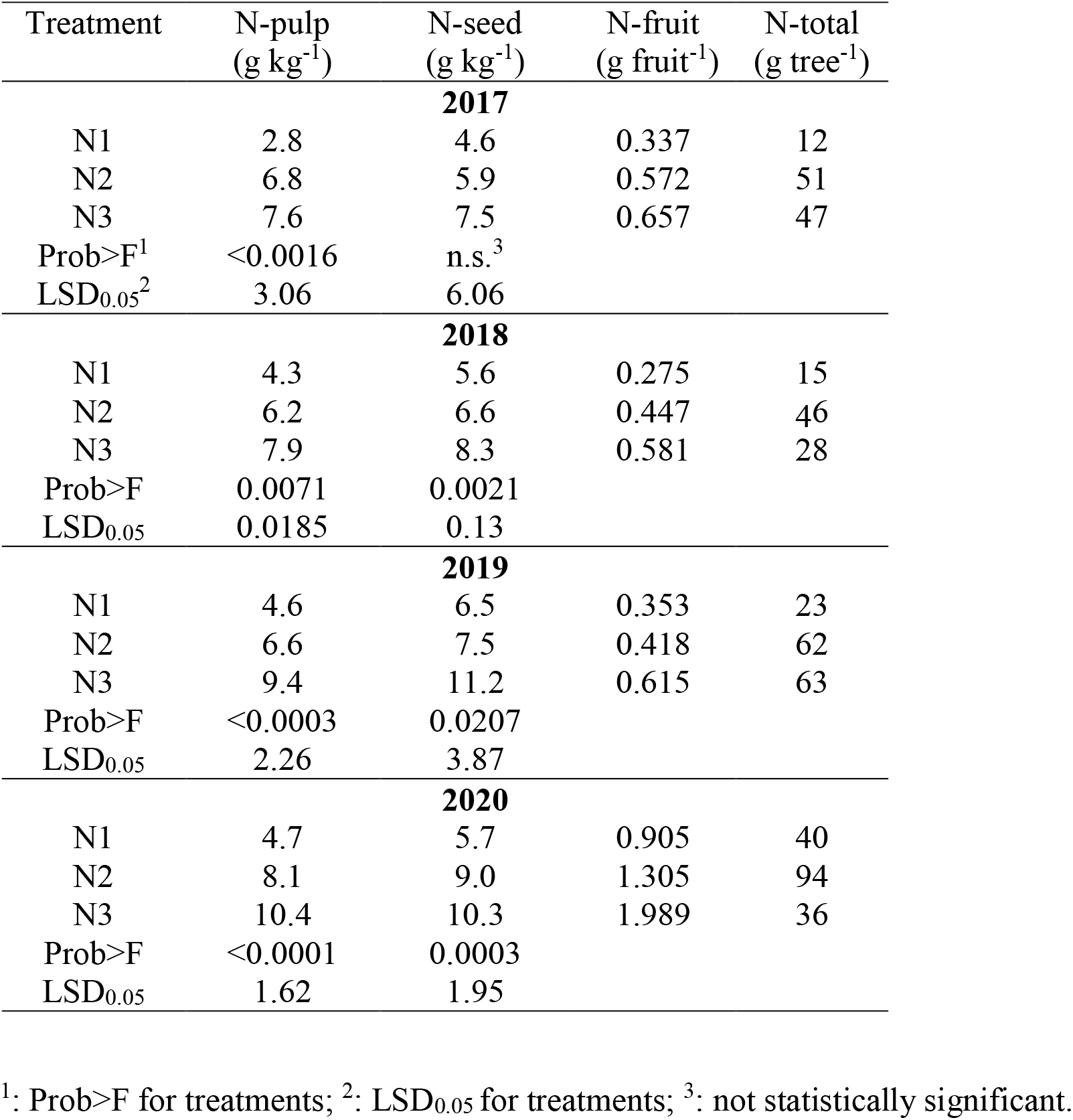
Nitrogen concentrations in fruit-pulp (N-pulp), fruit-seed (N-seed), or the entire fruit (including peel, pulp and seed). N-total is the total nitrogen quantity removed by fruit harvest for a tree.

### 3.3. Fruit color, TSS and acidity

The flavor of mango pulp is primarily determined by the ratio between soluble sugar and organic acid contents (Malundo et al., 2001; Léchaudel et al., 2005b). Citric acid, followed by malic acid, are the predominant acids (Léchaudel et al., 2005b), while sucrose, followed by fructose and glucose, are the major sugars in mango pulp (Medlicott and Thompson, 1985; Léchaudel et al., 2005b). TSS content in the present study significantly decreased as N-irg increased (Prob > F < 0.0001), while TA exhibited the opposite phenomenon (Prob > F < 0.0158) (Fig. 7). During ripening, the acid turns into sugar and therefore, N treatments may affect the rate of maturation. Note that the trend of TA decreases as N-irg increases (Fig. 7) and the opposite trend for An (Fig. 2), are in accordance with Léchaudel et al. (2005b) report that synthesis of citric acid is favored under conditions of slow fruit growth and low assimilation rate. Red skin color of mango fruit is a quality parameter, especially for export and it changes from green to yellow and to red during ripening. The degree of the skin red color significantly decreased as N-irg increased (Prob > F < 0.0001), in accord with TSS content and opposite with TA (Fig 7).

**Fig. 7.**
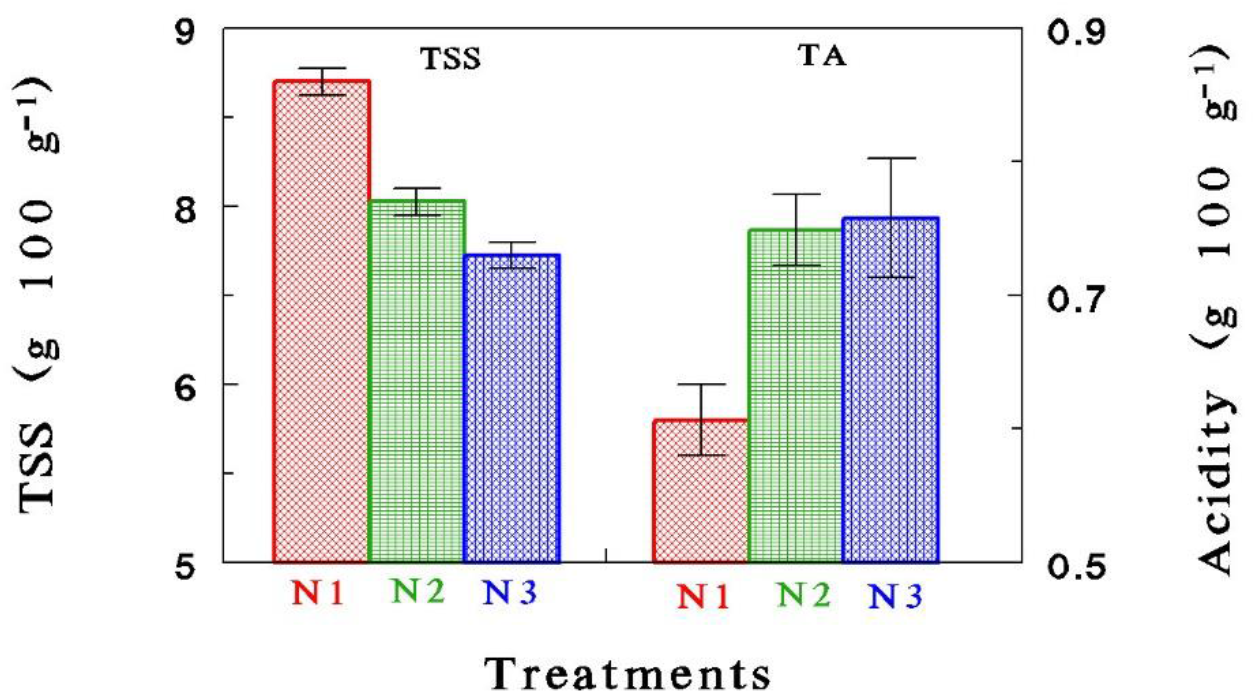
Total soluble sugars (TSS) and titrable acidity (TA) of mango pulp fruits in 2017. Vertical lines designate standard error values of four trees.

### 3.4. The nitrogen balance of mango trees

The results reveal mango’s nitrogen uptake and distribution. As previously suggested, the nitrogen uptake was correlated with the fruit production and accordingly it peaks during the reproductive period (Stassen and van Vuuren, 1997). We compared the calculated daily N quantities supplied by fertigation (N-irg) throughout 2019-20 growth season, to the calculated N-uptake (Eq. 1) and to the N quantities accumulated in the fruit (N-fruit). Total N quantities supplied throughout 2019-20 season to N1, N2 and N3 trees were 49(±7), 361(±42) and 691(±59) g tree^−1^, respectively, while annual N-uptake were 26(±5.2), 196(±18.4) and 185(±15.9) g tree^−1^, respectively. The N-efficiency ratio (N-uptake/N-irg) varied along the growth season in accordance with the seasonal variation of global solar radiation (Fig. 9), air temperature, vapor pressure deficit (VPD) and evapotranspiration (not presented). The lowest N-efficiency for the three treatments were measured on the middle of January (DOY=15, Fig. 9) and were: 0.41, 0.22 and 0.05 for N1, N2 and N3 trees, respectively. N-efficiency increased later as the climatic conditions became more favorable for plant development (global radiation is presented in Fig. 9). The averaged N-efficiency for the whole 2019-20 season were 0.52, 0.54 and 0.27, for N1, N2 and N3 trees, respectively. The low N-efficiency of N3 trees reflects excessive N fertilization. Fruits in the present study (Table 2, 2020 data) were the major N sinks, containing 40±4.9, 94±6.1 and 36±28.6 g N per tree, for N1, N2 and N3 trees, respectively. The accumulated N quantities in the 2020 fruits were: 0.82, 0.26 and 0.05 from the total N supplied and 1.54, 0.48 and 0.19 from N-uptake for N1, N2 and N3 trees, respectively). Note that the accumulated N quantities in the fruits of N1 trees exceed N1-uptake. Yet, fruit N quantities represents robust measured values, while N-uptake were calculated (eq.1) from data of N concentration analyses in solutions sampled every two weeks. It is highly possible that slight variations in N concentration in the irrigation and\or drainage solutions between two consecutive analyses induced the high standard error (Se) values of N-irg and N-uptk for N1 treatments. It is important to note that from 18 June (DOY 169) until harvest (30 August), the daily N quantities accumulated (N-fruit) in the N1-fruits were above the N quantities supplied via fertigation (N-irg) (Fig. 8).

**Fig. 8.**
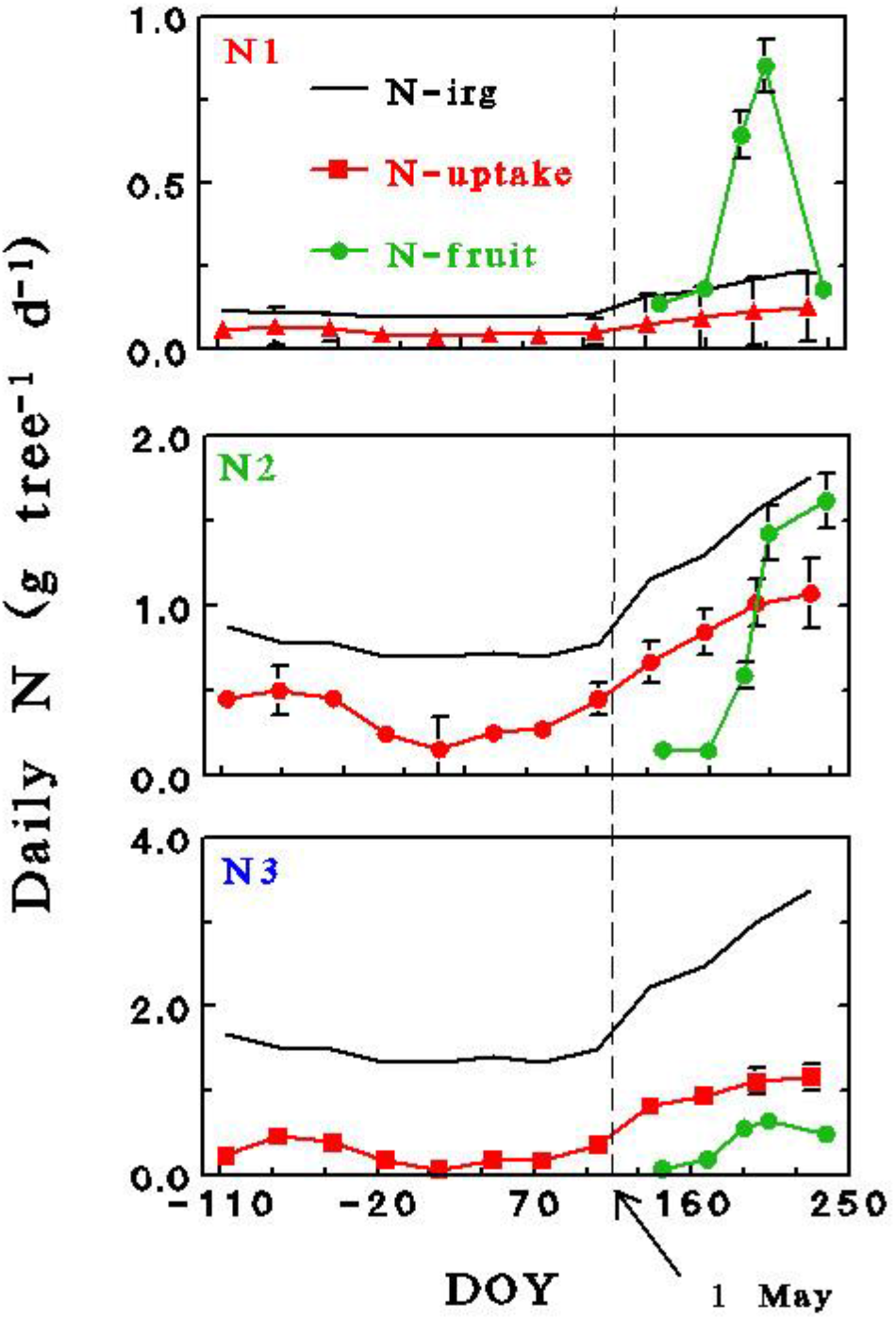
Effects of N treatments on the daily N quantities supplied every month by fertigation to (N-irg) throughout 2019-20 growth season, compared to the calculated N-uptake (Eq. 1) and to the N quantities accumulated in the fruit (N-fruit). Each value of N-fruit represents an average between the actual and the precedent point. Vertical solid lines designate standard error values, dashed vertical line designate the beginning of the reproductive season.

**Fig. 9.**
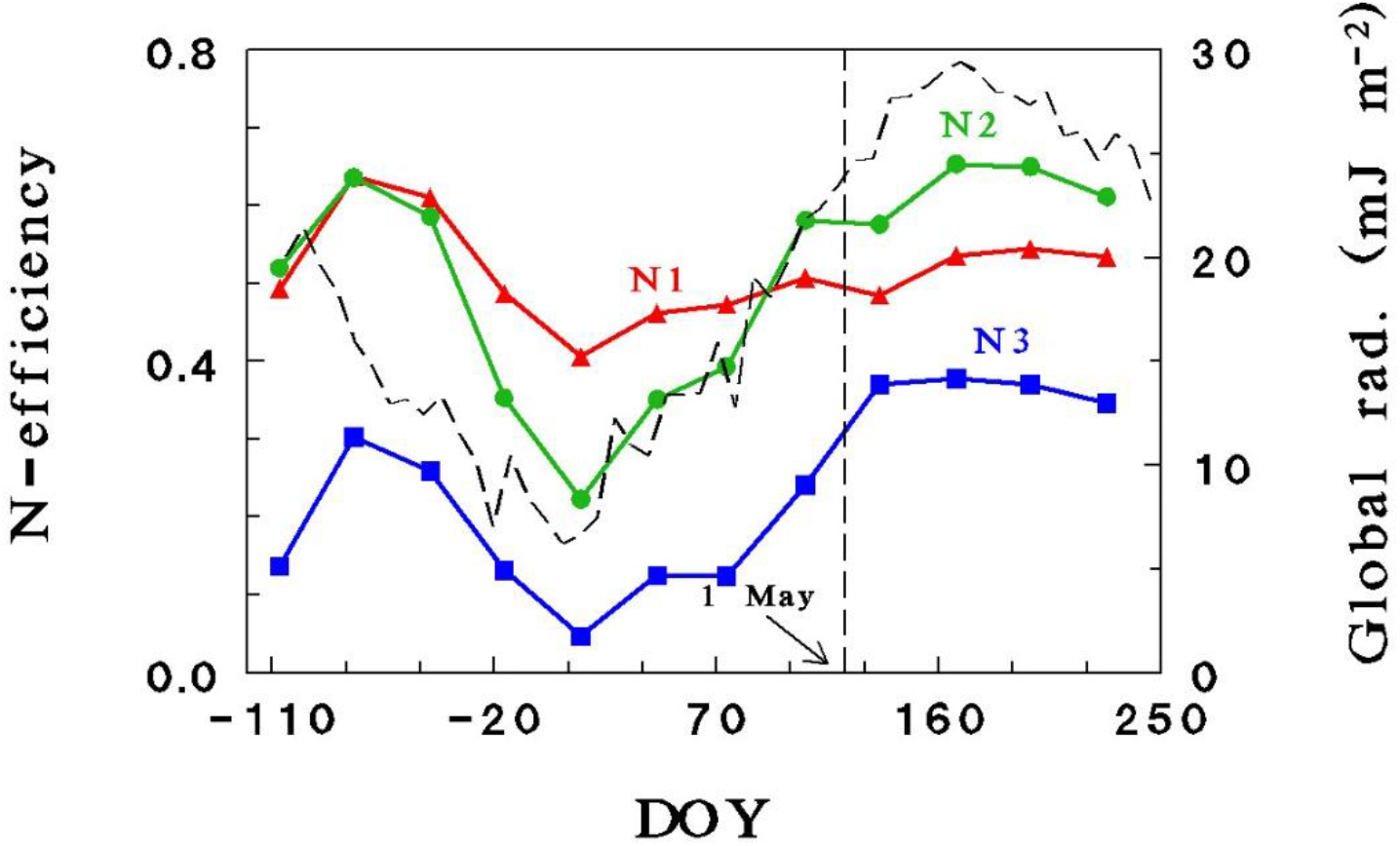
Variations of N-efficiency during 2019-20 season for N1, N2 and N3 trees compared with the global solar radiation (dashed line). Vertical dashed designate the beginning of the reproductive season.

The growing season was divided into two separate periods: vegetative (September 2019–April 2020) and reproductive (May–August 2020). Total leaf DW analyzed immediately after the 2020 harvest (30 August) was 2.3(±2.87), 7.3(±5.9) and 5.0(±4.2) kg/tree for N1, N2 and N3, respectively. To estimate the proportion of leaves on the tree at the end of the vegetative period (30 April 2020), we used the LAI_max_ measurements taken on 30 April and 30 August 2020. The proportions between the 30 April and 30 August 2020 measurements were: 0.59, 0.70 and 0.48 for N1, N2 and N3 treatments, respectively. We estimated that half of the leaves analyzed after harvest in 2020 were generated during the preceding year (before 2019). Consequently, the amount of N accumulated in the leaves during the vegetative or reproductive period was calculated as follows: total N in leaves (g/tree) analyzed after harvest in 2020 for each treatment x 0.5 (leaves produced before harvest, in 2019) x LAI_max_ proportion. Total N accumulation in leaves during the vegetative period were 5, 16 and 19 g tree^−1^ for N1, N2 and N3, respectively, and for the reproductive period were 7, 38 and 18 g tree^−1^ for N1, N2 and N3, respectively.

## 4. Discussion

### 4.1. The nitrogen balance of mango trees

The results reveal mango’s nitrogen uptake and distribution. As previously suggested, the nitrogen uptake was correlated with the fruit production and accordingly it peaked during the reproductive period (Stassen and van Vuuren, 1997). The low N-efficiency ratio of the vegetative period together with the high N-efficiency of the reproductive period indicate the involvement of an unaccounted-for N source or sink, or both. Roots, wood, bark, and new shoots accounted for more than a third of the total N stored in 6-year-old mango tree cv. Sensation (Stassen and van Vuuren, 1997; Stassen et al., 2000). Therefore, it is reasonable to assume that N quantities accumulated in these organs during the vegetative period of August–April could serve later throughout the reproductive period as N reserves for the high demand of the developing fruit. This notion is supported by the reduction in N concentration of leaves during fruit development (Figure 3). The importance of N-reserves for supplying the fruit demand complements those of Davie et al. (2000) and Léchaudel et al. (2005a) that starch reserves in the tree may supply an important part of the carbon needed by the fruit.

### 4.2. The low requirement of mango trees to N and the negative effect of excessive N

The negative effect of the high N concentration (N3) on productivity is common to most crops (e.g. Gashu et al., 2020), possibly due to straining of critical metabolic pathways (Sperling et al., 2019) or impairment of the reproductive: vegetative balance (Núñez-Elisea and Davenport, 1995). At the same time, the relatively low concentration that administered the damage was surprisingly low when compared to other fruit trees. Most fruit trees optimize productivity at 30 ppm N or higher (e.g. Erel et al., 2008; Sperling et al., 2019). The negative effect to yield of relatively low N concentration agrees with other mango studies which showed substantial reduction in yield when supplied with 23 ppm N (Torres et al., 2002) or 450 g N tree^−1^ year^−1^ (Nguyen et al., 2004). Also, the relatively high yield of the 2 ppm treatment compared with other crops that typically do not produce any fruits under such conditions (e.g. Sperling et al., 2019) are in agreement with other mango studies (Torres et al., 2002; Nguyen et al., 2004). The overall N uptake of N2 trees foe 2019-2020 season (196 g tree^−1^ year^−1^) is in accord with Stassen and Van Vuuren (1997) which estimated the uptake of 6 years old trees at 151 g tree^−1^ year^−1^. It should be noted that extrapolating the findings into other orchard must account for the planting density and the tree’s productivity. The low N requirements of mango’s are probably thanks to the low N content in the fruits (Figs 5, 6; Stassen and van Vuuren 1997).

N-irg concentration of 20 mg/L reduced fruit yield (Table 1, N3) and accordingly, water uptake (Table A.1) and gas-exchange characteristics (Fig. 2). Similarly, N addition to mango ‘Tommy Atkins’ trees induced flushes of new leaves and reduced floral induction (Núñez-Elisea and Davenport, 1995). Bud dormancy is a necessary step for floral initiation in mango (Davenport, 2009; Lu et al., 2012) and it is usually achieved by the application of water stress (Lu and Chacko, 2000; Carr, 2014) or cool temperatures (below 15 °C) (Núñez-Elisea and Davenport, 1995; Lu et al., 2012). Thus, excess N fertilization in the N3 treatment may have stimulated continuous vegetative growth, which impaired floral induction, or flower or fruit-set development, as reported recently for olive (Haberman et al., 2019) and almond (Sperling et al., 2019) trees. Obviously, it is difficult to distinguish cause from effect in this case, as alternatively, excess N might have caused fruit abscission that stimulated vegetative growth.

### 4.3. Impacts of high fruit:leaf ratio driven by N fertilization treatments on fruit maturation, gas exchange activity and P and K uptake

High fruit:leaf driven by low N-irg concentrations accelerated the fruit maturation rate as indicated by high TSS, low TA (Fig. 7) and the skin red color detailed in subchapter *3.4*. The sharp decreases of N-fruit value along the last two weeks before harvest (Fig. 8), also give an indirect support to the conclusion that high fruit:leaf ratio induces advanced maturation rate.

Deficient-N conditions resulted in high photosynthetic activity (Fig. 2), high leaf-P and K concentrations (Fig. 3) and high maturation rate as indicated by TSS:TA ratio and skin red color detailed in subchapter *3.4*. N deficiency together with adequate P nutrition has been shown to induce excess P uptake by algae (Chu et al., 2013, 2014, 2015); Zhu et al., 2015). A similar trend of increased P uptake under N-deficient conditions has also been reported for spring barley (Nátr et al., 1983), Tef (*Eragrostis tef* Trotter) (Gashu et al., 2020) and almond trees (Sperling et al., 2019). The mechanism by which high fruit:leaf ratio driven by N-deficient conditions induces high leaf-P and K concentrations in fruit trees is probably complex and may include several steps: (i) enhanced gas-exchange activities – a high fruit:leaf ratio induced a high demand for carbohydrates and subsequently, enhanced stomatal conductance and photosynthetic activity (Fig. 2 and Urban et al., 2004a,b); and (ii) stimulated P and K uptake – increased sink activity is associated with enhanced P redistribution. Hopkinson (1964) reported that carbohydrate translocation from leaves to other plant organs of cucumber (*Cucumis sativus*) is associated with high P import into the leaf. Therefore, it may be suggested that high P and K concentrations in mango leaves, and fruit may be related to the increase in fruit:leaf ratio driven by N deficiency. In addition, the low amino acid and nitrate contents require osmotic compensation, which comes from K, the major inorganic osmolyte in plants.

### 4.4. Tools for fertilization control

Although leaf-nutrient status is frequently used for fertilization control in the mango industry (Bally, 2009; Pinto et al., 2007), its efficiency for this purpose is questionable at least for N (Bally, 2009). Furthermore, the basic transformation from concentration value (gram N-P-K per gram DW) to quantity value (gram N-P-K per tree or hectare) is problematic because leaf and fruit weights vary during the growing season according to physiological and phenological aspects. Moreover, the common assumption that each nutrient’s concentration in the leaf is predominantly determined by its fertilization rate and therefore, may be considered an independent variable, is also questionable. In the present study, although P and K concentrations were kept constant in the irrigation water, N deficiency (N1) induced high leaf-P and K concentrations (Fig. 3) by the mechanisms detailed in section *4.1*.

The goal of fertilization should be to replace the nutrient quantities that have been removed by the fruit or re-stored for further tree development; accordingly, fertilization rate should be adjusted to coincide with the actual demand for each nutrient at each phenological stage. Taking into consideration nutrient quantities removed by the expected yield (Fig. 6) and the vegetative growth (Stassen et al., 2000), combined with fertilization efficiency coefficient will provide a solid foundation for any fertilization management program.

## 5. Conclusions

1. N fertilization of mango trees may be a double-edged sword, as slight differences in N-irg concentrations may generate N deficiency or excess, and significant differences in vegetative and reproductive yield.
2. The common assumption that leaf-nutrient concentration is predominantly determined by its fertilization rate and therefore, may be considered an independent variable, is questionable. Thus, the efficiency of leaf-nutrient analyses as a method for fertilization control in the mango industry is questionable.
3. N deficiency conditions generated low vegetative yield, high fruit:leaf ratio, high photosynthetic activity, high leaf-P and K concentrations, high sugar contents, and low acidity in the fruit. Excess N concentration stimulated continuous vegetative growth, impaired floral induction or flower and fruit-set development, or both, and reduced fruit yield.
4. Excess N concentration stimulated continuous vegetative growth, impaired floral induction or flower and fruit-set development, or both, and reduced fruit yield.
5. Fruit are the major N sinks in mango trees and accordingly, nitrogen uptake significantly increases during the reproductive period.
6. N quantities accumulated in roots, wood, bark, and new shoots throughout the vegetative period of August–April could serve later, during the reproductive period, as reserves for the developing fruit’s high demand for N.

## Abbreviations

An: assimilation rate
*g*_s_: stomatal conductance
DOY: day of year
DW: dry weight
EC: electrical conductivity
K-pulp: K concentration in the pulp
K-seed: K concentration in the seed
K-fruit: K quantities accumulated in a single fruit
K-total: totaled K quantities removed by the fruits
LAI_max_: maximal leaf area index
N-pulp: N concentration in the pulp
N-seed: N concentration in the seed
N-fruit: N quantities accumulated in a single fruit
N-total: totaled N quantities removed by the fruits
N-irg: N quantities supplied by fertigation
P-pulp: P concentration in the pulp
P-seed: P concentration in the seed
P-fruit: P quantities accumulated in a single fruit
P-total: totaled P quantities removed by the fruits
TA: titrable acidity
TSS: total soluble sugars

---

**Table A.1.**
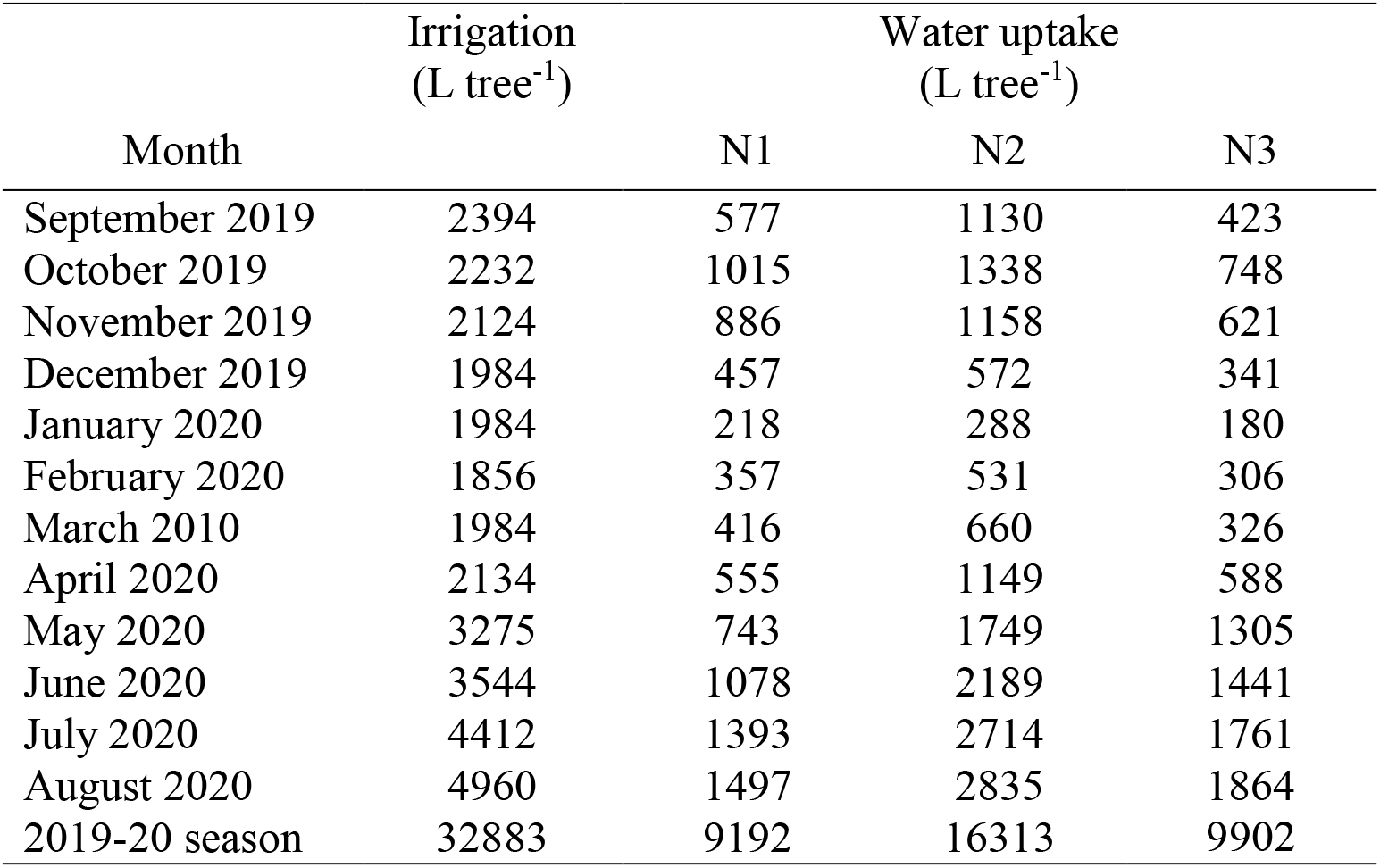
Monthly irrigation quantities supplied and water uptake of N1, N2 and N3 trees during 2019-20 season.

**Table A.2.**
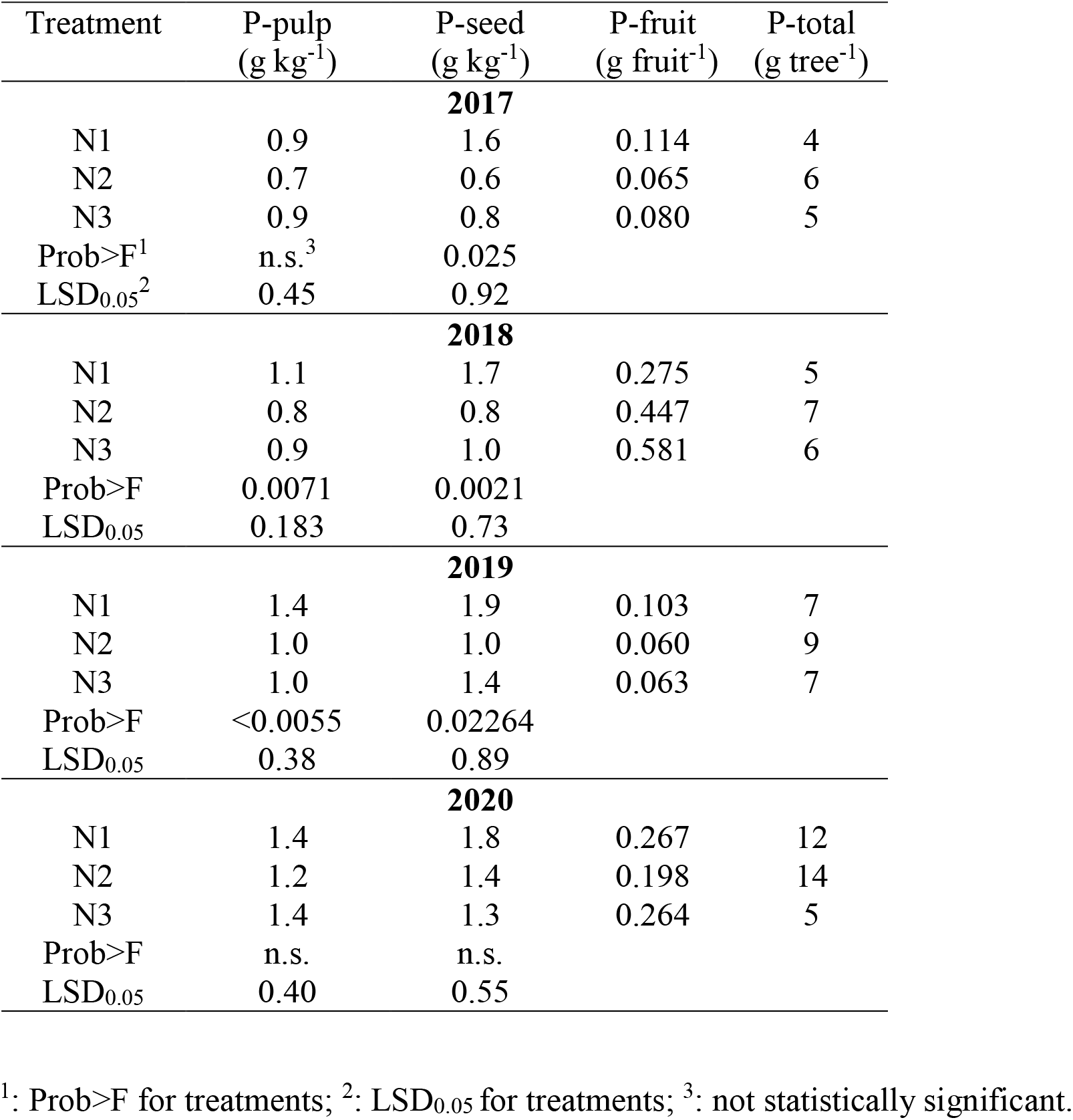
Phosphorus concentrations in fruit-pulp (P-pulp), fruit-seed (P-seed), or the entire fruit (including peel, pulp and seed). P-total is the total phosphorus quantity removed by fruit harvest for a tree.

**Table A.3.**
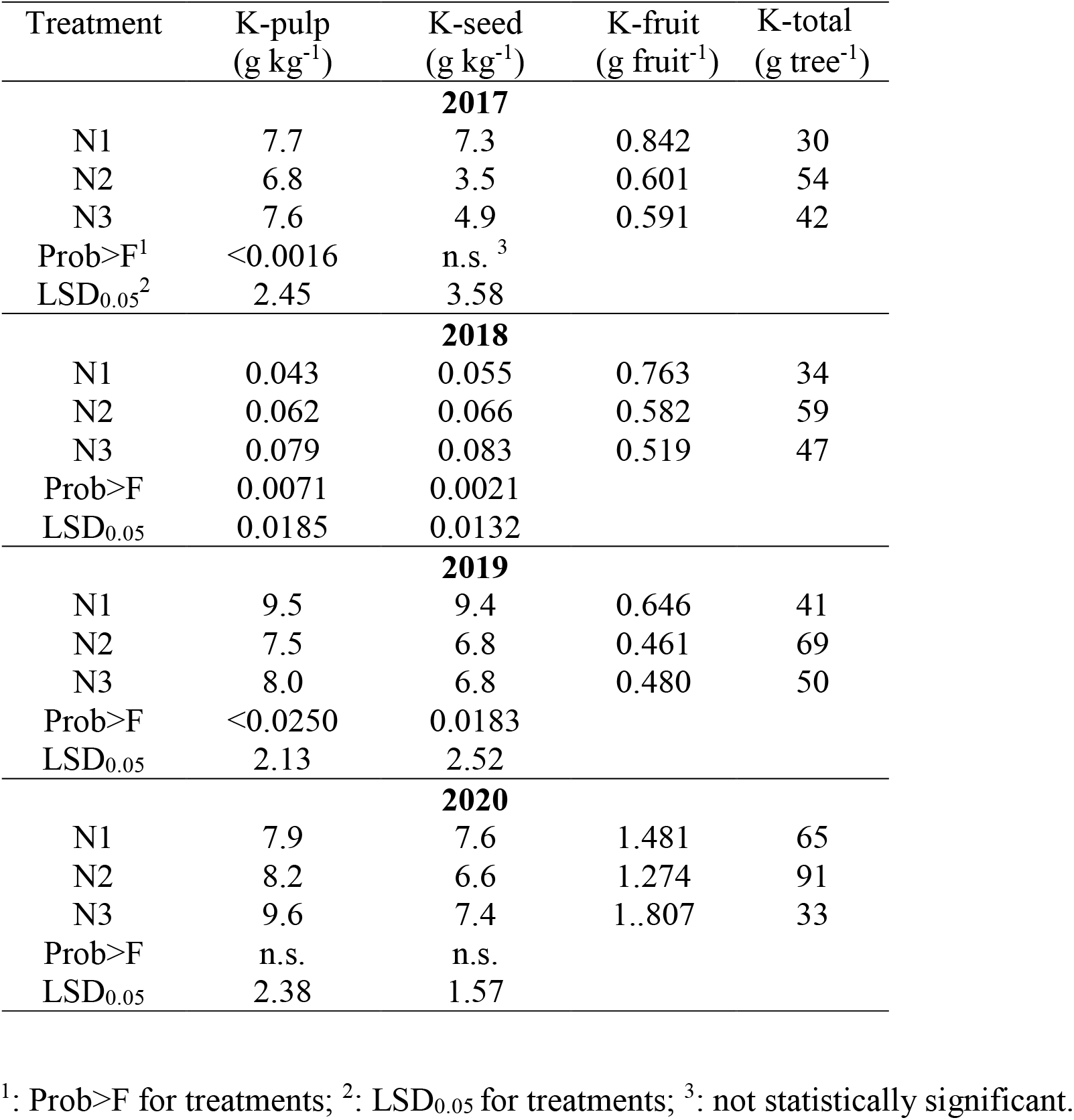
Potassium concentrations in fruit-pulp (K-pulp), fruit-seed (K-seed), or the entire fruit (including peel, pulp and seed). K-total is the total potassium quantity removed by fruit harvest for a tree.

